# Glomerulus-specific inhomogeneity of the basal activity map in the olfactory bulb supports olfactory-driven behavior

**DOI:** 10.1101/2025.11.24.690198

**Authors:** Stefan Fink, Natalie Fomin-Thunemann, Wen-Yu Tzeng, Katherine Figarella, Farzin Kamari, Yury Kovalchuk, Olga Garaschuk

## Abstract

Glomeruli are signal-processing units of the olfactory bulb (OB), playing a key role in many OB computations, including contrast enhancement, gain control, and odorant-selective habituation. In awake mice, we uncover an extremely stable inhomogeneous map of basal glomerulus-specific activity as a background against which olfactory signal processing is performed. This activity is strongly driven by (i) centrifugal cholinergic projections, (ii) endogenous, and (iii) airflow-evoked spiking of olfactory sensory neurons and, to a small extent, (iv) by the odor environment. Importantly, early olfactory impairment in a mouse model of Alzheimer’s disease is parallelled by the loss of glomerular map inhomogeneity and diminished cholinergic innervation. These results reveal an important layer in the signal-processing network of the OB, likely acting by increasing the variance in and dynamic range of the system via glomerulus-specific functional inhomogeneity.

**Teaser:** A new layer in the signal-processing network of the olfactory bulb supports olfactory-driven behavior.

## Introduction

The olfactory bulb glomeruli are fundamental units representing the first stage of sensory processing in olfaction. In mice, a typical glomerulus receives inputs from ∼1000 olfactory sensory neurons (OSNs), all expressing the same type of odor receptor (OR). Within the glomerulus, the OSN axons synapse on primary dendrites of 20-50 mitral/tufted (M/T) cells and hundreds of juxtaglomerular interneurons (*1, 2*). Experimental and computational modeling data suggest that intraglomerular computations play a key role in contrast enhancement, gain control, odorant-selective habituation, and odorant-specific suppression of M/T cell outputs (*3–5*). Because the chemoreceptive properties of different OSNs overlap such that each odor activates many ORs to a greater or lesser extent, it is largely assumed that it is the combinatorial pattern across the set of corresponding odor-activated glomeruli that forms the basis for odor perception and identification. Accordingly, perceptually similar odors activate a correspondingly greater number of common ORs/glomeruli, thus evoking highly overlapping primary odor representations, which are then decorrelated by the OB circuitry (*6, 7*). This logic, however, implicitly assumes that odor-evoked OSN signals are projected onto a layer of “resting” glomeruli, with some nonspecific background noise (*8–11*). This noise might comprise spontaneous activity of OSNs, known to fire at frequencies between 0 and 7 Hz (*12–16*), as well as that of spontaneously active juxtaglomerular interneurons and M/T cells (*17, 18*), randomly distributed in the glomerular layer in a salt-and-pepper fashion. Here, by using *in vivo* two-photon Ca^2+^ imaging in awake head-restrained mice, with neuronal expression of a ratiometric Förster resonance energy transfer (FRET)-based Ca^2+^ indicator Twitch-2B under the chronic cranial window, we challenge the above view by showing that the majority of glomeruli are not silent. Rather, the “resting state” of the glomerular layer of the OB is characterized by highly inhomogeneous, stable, and glomerulus-specific patterns of endogenous activity. Moreover, the early disease-related loss of this resting state inhomogeneity impacts olfactory-driven behavior.

## Results

### Inhomogeneity of the glomerular map in resting awake animals

All experiments were conducted in awake head-restrained mice, which were extensively trained and habituated to the imaging setup before recordings. As shown in Fig. 1A, structures with different background-corrected ratios of the acceptor to donor fluorophore fluorescence (cpVenus^CD^/mCerulean3, hereafter referred to as Twitch-2B ratios) coincided in shape and size with individual glomeruli, visible on the fluorescence image of the glomerular layer (left panel).

**Fig. 1.**
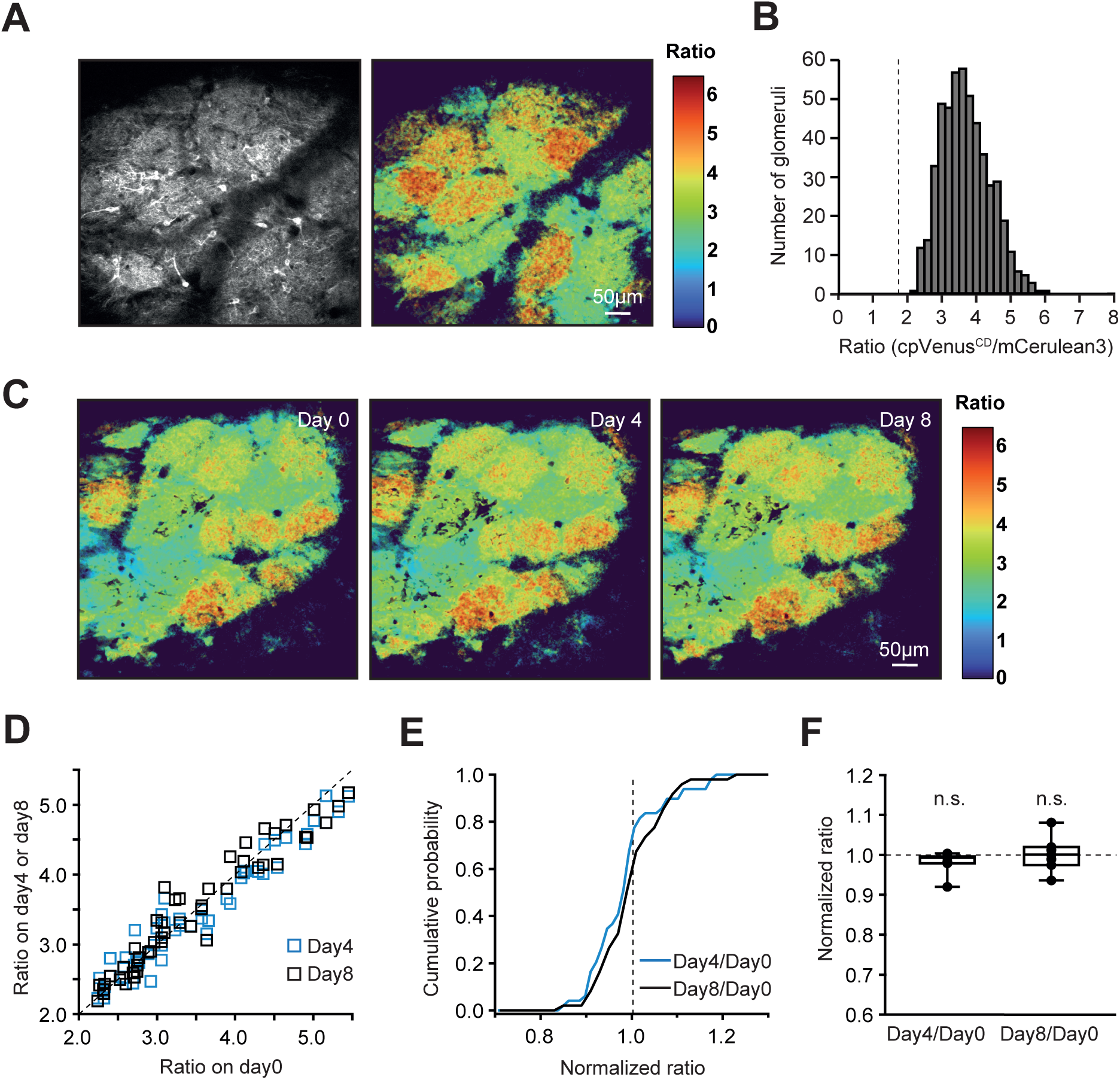
Inhomogeneity and stability of glomerular basal Twitch-2B ratios in awake mice. (A) Left: average intensity projection image of the dorsal surface of the OB (40-58 µm depth), labeled with the Ca2+-indicator Twitch-2B. Right: the corresponding average projection map of the color-coded cpVenusCD/mCerulean3 ratios, hereafter referred to as “color map”. Note that each glomerulus appears on the map in a distinct, largely homogeneous color. (B) Histogram illustrating the distribution of basal Twitch-2B ratios of individual glomeruli (n=506 glomeruli, 23 mice). A broken vertical line marks the spiking threshold (see fig. S2C). (C) Sample glomerular color maps of an OB region taken at three different time points (Day 0, Day 4, and Day 8). (D) Scatter plot showing basal glomerular Twitch-2B ratios recorded at Days 4 and 8 plotted against the corresponding ratios recorded at Day 0 (n=49 glomeruli, 7 mice). Diagonal broken line: the unity line. (E) Cumulative probability plot displaying the distribution of normalized basal glomerular Twitch-2B ratios in the data set shown in D. (F) Box plot showing the medians (per mouse) of normalized ratios, which are not significantly (n.s.) different from one (Friedman’s test P=0.18, χ2(2)=3.4).

The mean basal ratios recorded from different glomeruli ranged from 2.16 to 5.92 (Fig. 1B, n=506 glomeruli in 23 mice), well within the dynamic range of the Twitch-2B Ca^2+^ sensor, ranging from 1.3 (see below) to 8.5 (*19*). The observed inhomogeneous maps were surprisingly stable across different experimental days (Fig. 1, C to F and fig. S1). To test whether the differences in the basal Twitch-2B ratios between individual glomeruli reflect differences in action potential firing of underlying network elements, we applied a sodium channel blocker tetrodotoxin (TTX, 5 µM) on top of the cranial window with a slit (*20*). Topical TTX application strongly reduced the basal Twitch-2B ratios (below 1.8), completely wiping out map inhomogeneity (fig. S2). We concluded, therefore, that basal Twitch-2B ratios of different glomeruli reflect spiking frequencies of contributing neuronal elements and that glomeruli with basal Twitch-2B ratios ≤ 1.8 can be considered non-spiking, whereas the ratios ≥ 2 reflect spiking of contributing neuronal elements (fig. S2C). This conclusion was further substantiated by recordings at a higher sampling rate (7.76 Hz), which revealed ongoing fluctuations of the intracellular Ca^2+^ levels in glomeruli with high Twitch-2B ratios (fig. S3). Surprisingly, neuronal elements belonging to one glomerulus seemed to have rather similar colors (i.e., Twitch-2B ratios) and thus spiking frequencies, underscoring the functional unity of individual glomeruli(*18*). According to our cell soma-based *in situ* calibration (*19*), ratio values of 2-3 correspond to mean neuronal spiking frequencies of 0-5 Hz, whereas ratio values of 6 - to those of 20 Hz. Of note, in awake mice, we have not encountered any non-spiking glomeruli (Fig. 1B).

### Underlying cellular and molecular mechanisms

Next, we focused on mechanisms sustaining the glomerular map inhomogeneity in the resting state. To estimate the contribution of the OSNs, we applied 50 µM TTX in 10 µl PBS intranasally, re-examined the glomerular activity map 20-30 min after the TTX application (Fig. 2, A to B), and quantified the blocker’s effect size (Fig. 2C, 38.02±13.52, n=69 glomeruli, 6 mice; see Materials and methods). This treatment resulted in a significant reduction of the basal Twitch-2B ratios of individual glomeruli, accompanied by a decrease in the apparent map inhomogeneity. It also strongly inhibited the odor-evoked glomerular signals (fig. S4). The basal Twitch-2B ratios of individual glomeruli and the apparent map inhomogeneity largely recovered 24 hours after the treatment (Fig. 2, A to C). The application of 10 µl PBS alone also reduced basal Twitch-2B ratios, albeit to a smaller extent (median effect size 11.96±11.75, n=33 glomeruli, 3 mice), likely due to a change in the airflow through the nose (see the description of Fig. 3 below).

**Fig. 2.**
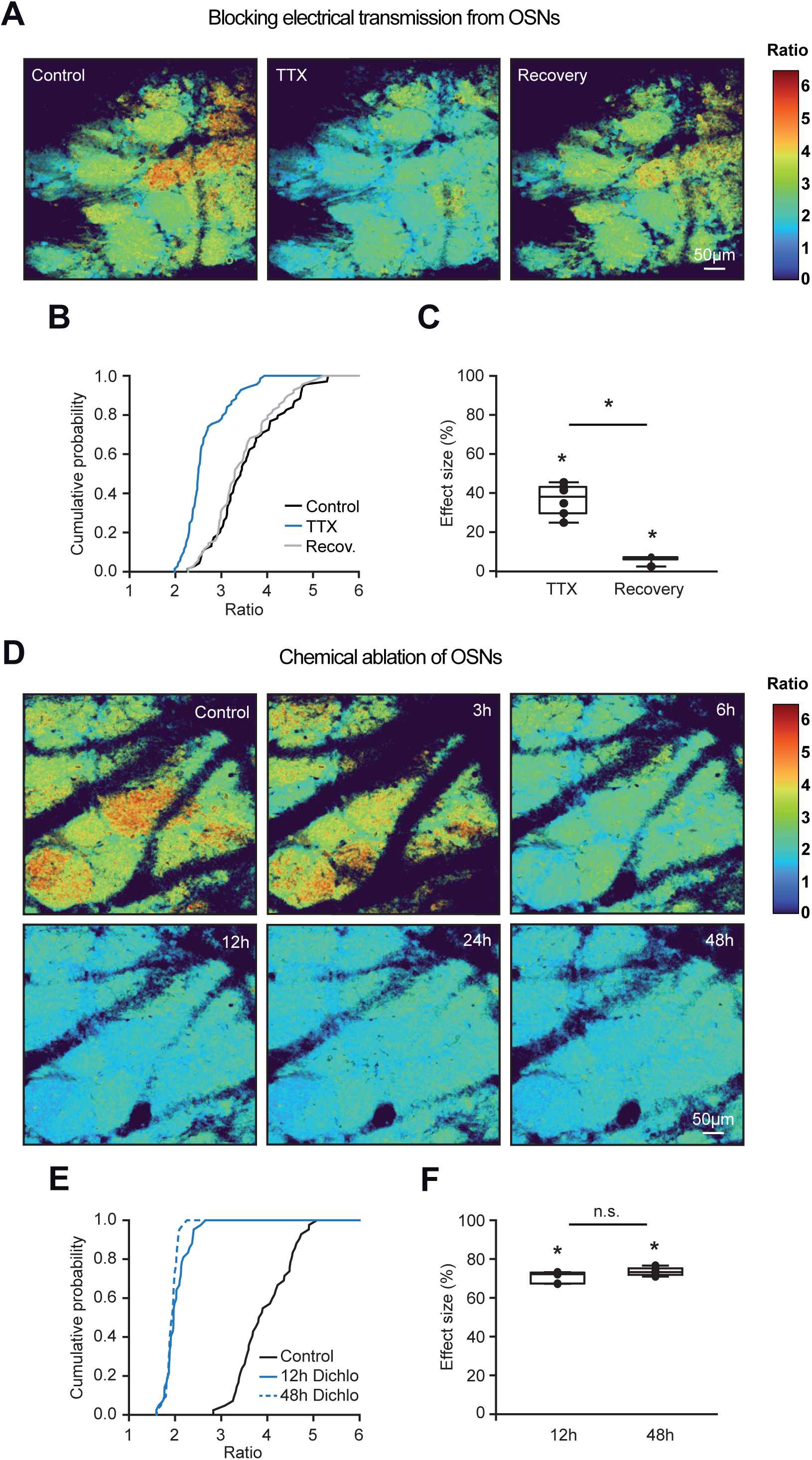
Activity maps depend on action potential firing of intact OSNs. (**A**) Color maps illustrating a reduction of basal Twitch-2B ratios upon intranasal application of 50 µM tetrodotoxin (TTX) and their partial recovery 24 hours after the application. (**B**) Cumulative probability plot showing the distributions of basal Twitch-2B ratios before (black), during (blue), and 24 hours after (gray) TTX application (n=70 glomeruli, 6 mice). (**C**) Box plot displaying the median (per mouse) effect sizes (calculated as described in Materials and methods; n=6 mice) of TTX blockade and recovery. TTX and recovery effect sizes significantly differ from zero (Friedman’s test P=2.5*10^-3^, χ^2^(2)=12, post-hoc Conover test with Bonferroni correction: P<2.2*10^-16^ for all comparisons). (**D**) Color maps illustrating the gradual decrease of basal glomerular Twitch-2B ratios within hours after i.p. injection of dichlobenil (50 µg/g body weight). (**E**) Cumulative probability plot showing distributions of basal ratios before (black), 12 (blue), and 48 hours (broken blue line) after dichlobenil injection (n=42 glomeruli, 6 mice). (**F**) Box plot showing the effect sizes of induced changes 12 and 48 hours after dichlobenil injection (Friedman’s test, P=5.7*10^-3^ χ^2^(2)=10.3; post-hoc Conover test with Bonferroni correction: P=1.7*10^-3^ for 12h, P=4.5*10^-5^ for 48h, and P=5.3*10^-2^ for comparison between 12 and 48 hours).

**Fig. 3.**
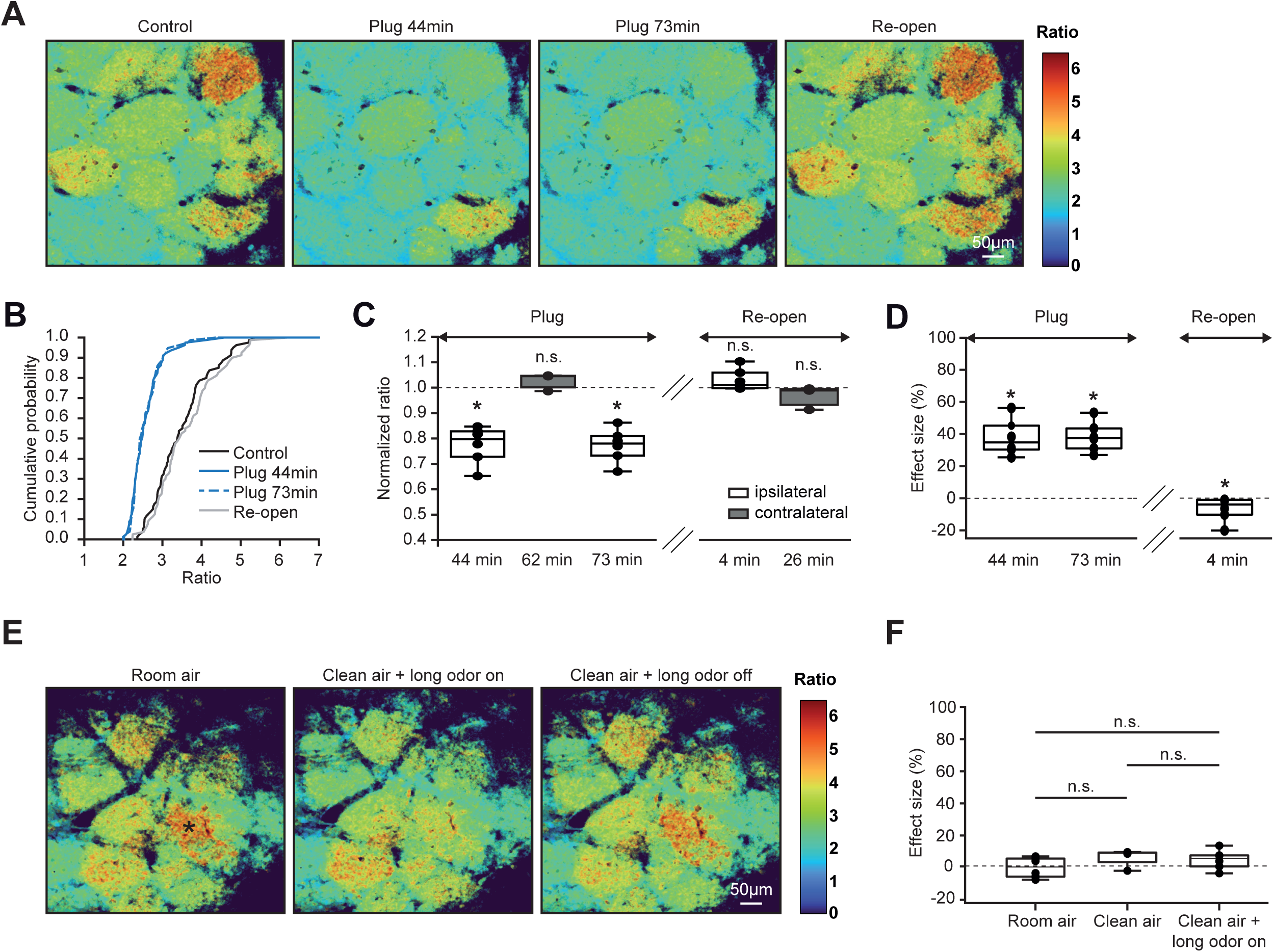
Activity maps depend on the nasal airflow. (**A**) Color maps illustrating basal Twitch-2B ratios before (Control), during (Plug), and after (Re-open) reversible plugging of one nostril. (**B**) Cumulative probability plot showing distributions of basal ratios before (black), during (blue), and after (gray) the nostril plugging (n=80 glomeruli, 6 mice). (**C**) Box plot showing the medians (per mouse) of normalized (to immediately preceding control) basal ratios of glomeruli ipsi- (white, n=6 mice) and contralateral (gray, n=3 mice) to the plugged nostril. Time points on the x-axis indicate the duration of nostril plugging (Plug) or time after plug removal (Re-open; mixed model ANOVA with post-hoc Tukey test, P<10^-4^ for 44 and 73 min and P ≥ 0.66 for all other comparisons). (**D**) Box plot showing effect sizes of ratio changes (Friedman’s test, P=1.03*10^-3^ χ^2^(3)=16.2, post-hoc Conover test with Bonferroni correction: compared to zero effect size P=2.1*10^-4^ for both 44 and 73 min, and P=9*10^-3^ for re-open). (**E**) Color maps illustrating basal Twitch-2B ratios in Room air, during continuous application of isoamyl acetate (IAA, 25-29 min after Odor on, 0.18% s.v.) and 10-14 min after the application (Odor off). (**F**) Box plot showing effect sizes of ratio changes measured during the 30-min-long period in Room air, Clean air, and Clean air + long odor on (Kruskal-Wallis test, P=0.13; n=6, 4, 6 mice, respectively).

As the data described above likely reflected only a partial blockade of the OSN Na^+^ channels (due to blocker release far from the OSNs or its impeded penetration through the mucus), in the next series of experiments, we chemically ablated OSNs using olfactotoxin dichlobenil (*21*). Already 6h after the i.p. injection of dichlobenil, we observed a visible reduction in the apparent map inhomogeneity, with a strong and significant blockade of glomerular activity seen between 12 and 48 h after the drug application (Fig. 2, D to F). Together, these data identify action potential-driven OSN activity as a strong contributor to the glomerular map inhomogeneity in the resting state.

The action potential (AP) firing in OSNs can occur spontaneously, in a cell-autonomous fashion, or can be driven by either the ambient odorants or the airflow. To distinguish between these possibilities, we first reversibly plugged one nostril with the silicon elastomer Kwik-Cast (Fig. 3, A to D, see Materials and methods). Because in mice odors are inhaled through the nostrils into two segregated nasal passages (*22, 23*), some 40-80 min after nostril plugging, the ipsilateral hemibulbs showed a significant and reversible reduction of the basal Twitch-2B ratios (normalized to control median (per mouse) values: 0.78 ± 0.1, n=6 mice). Despite the reduction, none of the glomeruli became silent (basal Twitch-2B ratios of all glomeruli ≥ 2, Fig. 3B), likely due to the endogenous firing of OSNs. The basal Twitch-2B ratios of glomeruli in contralateral hemibulbs did not change (Fig. 3C). Similar results were obtained when calculating the effect size of the nostril blockade (Fig. 3D).

To estimate the contribution of ambient odorants to map inhomogeneity, we first established the background map variability by monitoring the basal glomerular Twitch-2B ratios for 30 minutes in room air, then switched for 30 s to clean air, applied from an air tank and devoid of ambient odorants, and subsequently added a small concentration of the odorant to the clean air (0.18% of saturated vapor (s.v.), 30-min-long application) (Fig. 3, E to F). In contrast to the nostril blockade (Fig. 3D), the effect sizes of all aforementioned manipulations did not differ from 0 and were similar to the effect size of room air (Fig. 3F).

We noticed, however, that some glomeruli (e.g., the one marked by an asterisk in Fig. 3E) either increased or decreased their Twitch-2B ratios during the aforementioned manipulations, reflecting heterogeneous changes that were masked at the group level. To capture these subtle but meaningful variations, we reanalyzed our dataset using the absolute ratio changes (ΔR). The fluctuations of the basal glomerular Twitch-2B ratios were negligible in room air. The ratios did not change significantly in clean air but did show a small but significant change during the long odorant application (fig. S5). This suggests a small but measurable contribution of ambient odorants to the AP firing of OSNs.

To estimate whether the centrifugal inputs, arising from the subcortical nuclei (*24, 25*), might contribute to the glomerular activity map, we first studied the effect of different anesthetics (Fig. 4, A to D), collectively known to modulate the ascending reticular activating system (ARAS) (*26, 27*). The experiments were conducted every second day, with only one anesthesia regimen tested on that day, and corresponding awake recordings taking place right before the anesthesia induction. Isoflurane, ketamine/xylazine or MMF (medetomidine, midazolam, fentanyl) significantly blocked the glomerular activity (Fig. 4, A to D). The strongest effect was caused by isoflurane (median effect size: 62.25 ± 4%, n=6 mice).

**Fig. 4.**
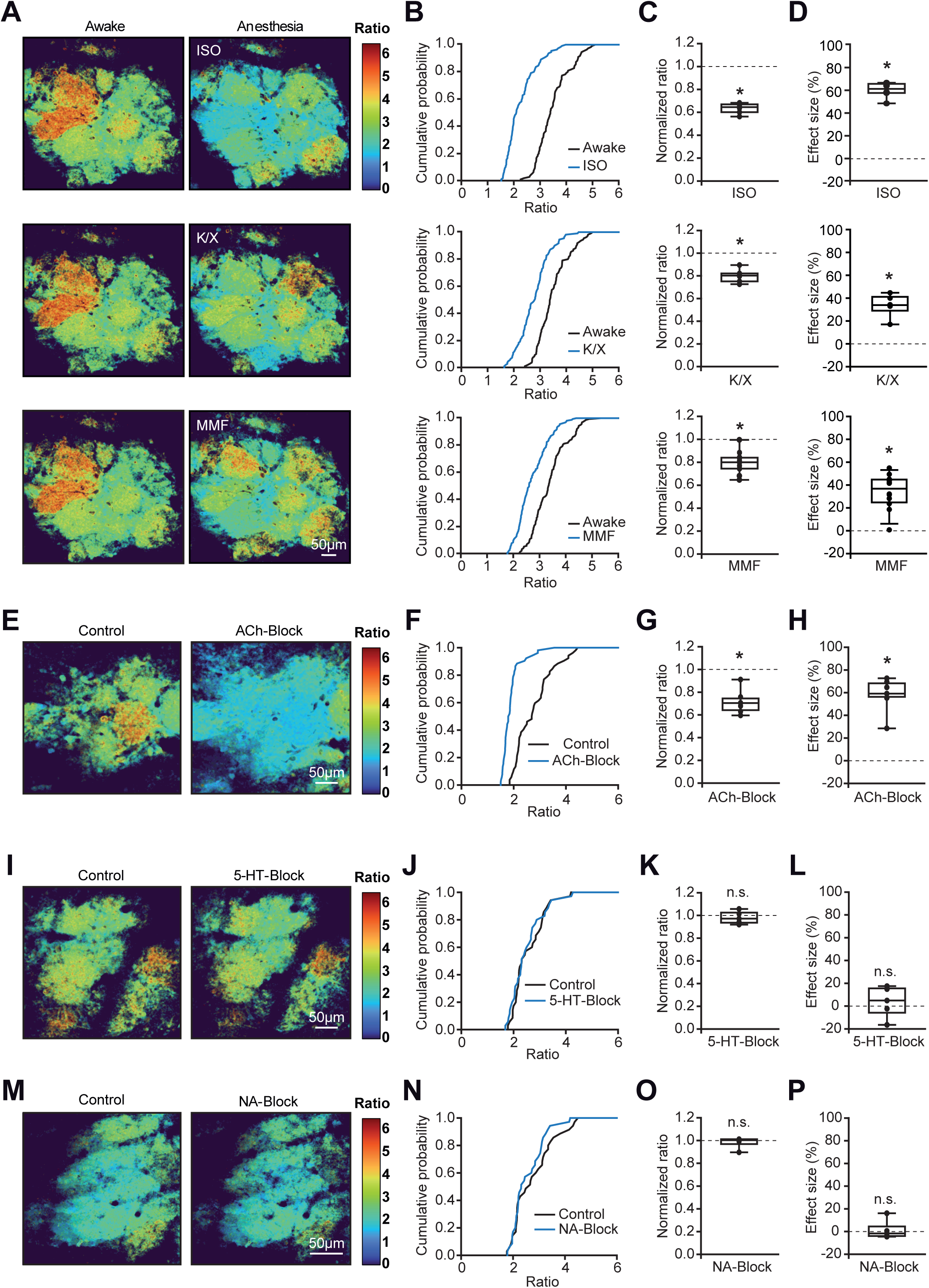
Modulation of activity maps by anesthesia and centrifugal inputs. (**A**) Color maps, acquired pairwise (awake and anesthesia) from the same animals every second day and illustrating the effect of isoflurane (ISO, top), ketamine/xylazine (K/X, middle), and medetomidine/midazolam/fentanyl (MMF, bottom) on the basal Twitch-2B ratios. (**B**) Cumulative probability plots showing distributions of basal ratios under one of the 3 anesthetic regimens (ISO, K/X, MMF, blue) or the respective awake state (black). ISO and K/X: n=110 glomeruli, 6 mice; MMF: n=152 glomeruli, 12 mice. (**C**, **D**) Box plots illustrating the median (per mouse) normalized ratios (C) and effect sizes (D) for the 3 different types of anesthesia (one sample t-test, ISO: P=6.3*10^-6^ and 3.8*10^-6^; K/X: P=4.2*10^-4^ and 3.1*10^-4^; MMF: P=2.7*10^-6^ and 7.6*10^-6^, respectively). (**E**, **I**, **M**) Color maps illustrating basal Twitch-2B ratios during topical application of HEPES-buffered Ringer solution in the absence (Control) and the presence (**E**, ACh-Block) of cholinergic receptor blockers mecamylamine and scopolamine; (**I**, 5-HT Block) of serotonergic receptor blocker methysergide; and (**M**, NA-Block) α1-adrenergic receptor blocker prazosin. (**F, J, N**) Cumulative probability plots showing distributions of basal Twitch-2B ratios in control and during one of the following pharmacological treatments: (**F**) mecamylamine (115 µM)/scopolamine (50 µM), n=52 glomeruli, 7 mice; (**J**) methysergide (4 mM), n=35 glomeruli, 5 mice; (**N**) prazosin (100 µM), n=35 glomeruli, 5 mice. (**G, K, O**) Box plots showing the median (per mouse) normalized ratio values for the 3 pharmacological treatments. One-sample t-tests: (**G**) P=3.13*10^-4^, n=7 mice; (**K**) P=0.55, n=5 mice. (**O**) Wilcoxon Signed-Rank test: P=0.81, n=5 mice. (**H, L, P**) Box plots showing the respective median (per mouse) effect sizes. One-sample t-tests: (**H**) P=4.1*10^-5^, n=7 mice; (**L**) P=0.59, n=5 mice. (**P**) Wilcoxon Signed-Rank test: P=0.81, n=5 mice.

As, in addition to modulating ARAS, anesthesia also impacts respiration, we next tested the specific contribution of cholinergic, noradrenergic, and serotonergic projections by applying a saturating dose of the respective receptor channel blockers on top of the cranial window with a slit (Fig. 4, E to P). The application of nicotinergic and muscarinergic receptor blockers, mecamylamine (115 µM) and scopolamine (50 µM), strongly and significantly inhibited glomerular activity (Fig. 4, E to H), with 50% of glomeruli becoming silent (basal Twitch-2B ratios < 1.8), whereas the blockers of serotonergic (5-HT, 4 mM methysergide, Fig. 4, I to L) and noradrenergic (NA, 100 µM prazosin, Fig. 4, M to P) receptors had no effect. Together, these experiments identify cholinergic projections from the basal forebrain (*25*) as a strong contributor to the map inhomogeneity in the glomerular layer of the main olfactory bulb.

### Role of the glomerular map inhomogeneity for early olfactory impairment in AD

To test the functional significance of the basal glomerular activity map, we chose Alzheimer’s disease (AD), known both for the early impairment of olfaction and the early degeneration of the brain-wide cholinergic projections (*28, 29*). Of note, these pathologies are reproduced in mouse models of the disease (*30, 31*). Here, we used the APPswe/PS1G384A mice (hereinafter referred to as AD mice, see Materials and methods), showing early neuronal hyperactivity in the cortex and hippocampus as well as impaired spatial and working memory at the age of 6-8 months (*32, 33*). In the cortex of these mice, amyloid β (Aβ) plaques start forming at ∼3 months of age (*32*). However, the olfactory bulbs of 3-month-old mice (n=4) were plaque-free, and in 5-7-month-old mice (n=5) plaques were seen only in the granule cell layer but not in the glomerular or external plexiform layers (fig. S6).

Surprisingly, the basal Twitch-2B ratios of individual glomeruli in AD mice were lower than in WT mice already at 3-4 months of age (Kolmogorov–Smirnov test, P=9.16*10^-11^) and remained at the reduced level thereafter (Fig. 5, A to C, G to I). Moreover, the inhomogeneity of the glomerular activity map, assessed as the inter-glomerular variance of the basal Twitch-2B ratios (see Materials and methods), also decreased in AD compared to WT mice as early as 3-4 months of age (Fig. 5D, here and below, weighted Wilcoxon-Mann-Whitney test with Bonferroni correction, P=1.6*10^-4^). This difference remained significant also at 5-7 months of age (Fig. 5J, P=2.8*10^-4^). The odor detection ability of AD mice, measured in a buried food pellet test (*34*), was also significantly impaired as early as 3-4 months of age (but not in 1-month-old mice (Fig. 6C)) and worsened further with disease progression (Fig. 5, E and K). In contrast, the sensitivity of odor-evoked responses (estimated from EC_50_ values of the respective dose-response curves) was still normal in 3-4-month-old AD mice (Fig. 5F) but deteriorated till 5-7 months of age (Fig. 5L). We did not observe any significant correlation between the odor sensitivity (EC_50_ values) of individual glomeruli and their basal Twitch-2B ratios (Fig. 5, M to N).

**Fig. 5.**
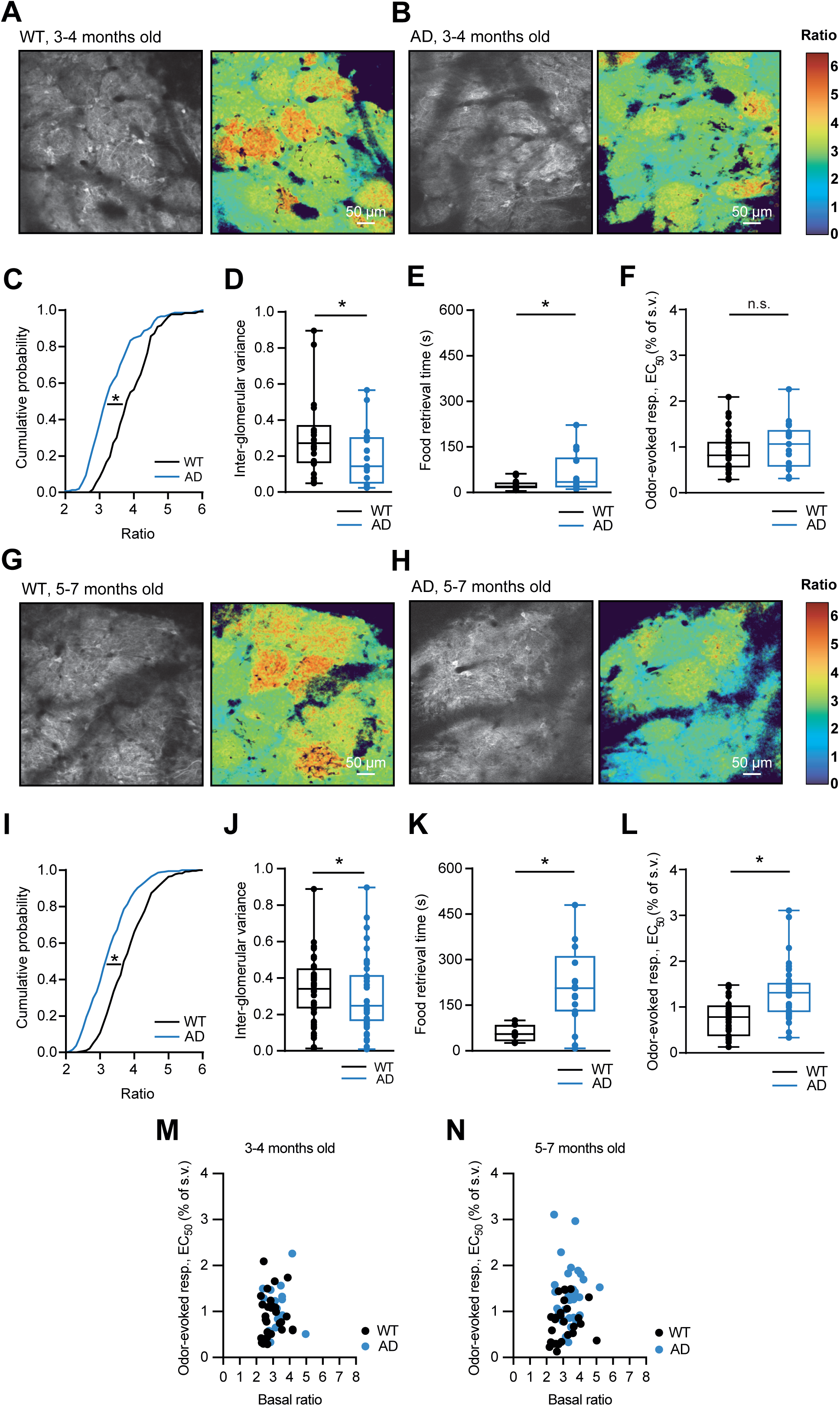
Reduced map inhomogeneity accompanies impaired sense of smell in a mouse model of AD. (**A, B, G, H**) Sample average intensity projection images (∼40-60 µm depth) of the dorsal surface of the OB labeled with the Ca^2+^-indicator Twitch-2B (left) and the corresponding color maps (right) taken in WT and AD mice of respective ages. (**C, I**) Cumulative probability plots illustrating distributions of basal Twitch-2B ratios in WT and AD mice of respective ages (Kolmogorov-Smirnov test, 3-4 months: P=9.16*10^-11^, n=133 glomeruli from 4 WT and 150 glomeruli from 4 AD mice; 5-7 months P=1.06*10^-15^, n=278 glomeruli from 8 WT and 380 glomeruli from 11 AD mice). (**D**, **J**) Box plots illustrating the interglomerular variances (calculated as described in Materials and methods) in WT and AD mice of respective ages (exact two-sided weighted Mann-Whitney-Wilcoxon test, 3-4 months: P=1.6*10^-4^, n=15 WT and 14 AD mice; 5-7 months: P=2.8*10^-4^, n=6 WT and 17 AD mice). (**E, K**) Box plots illustrating the time spent retrieving the chocolate in a buried food pellet test by WT and AD mice of different ages (Wilcoxon-Mann-Whitney test, 3-4 months: P=0.02, n=15 WT and 14 AD mice; 5-7 months: P=3*10^-4^, n=6 WT and 17 AD mice). (**F**, **L**) Box plots illustrating the EC_50_ values of ETI dose-response curves (see Materials and methods) in WT and AD mice of respective ages (3-4 months: t-test, P=0.38, n=29 glomeruli from 5 WT and 17 glomeruli from 4 AD mice; 5-7 months: Wilcoxon-Mann-Whitney test, P=3*10^-4^, n=27 glomeruli from 7 WT and 37 glomeruli from 6 AD mice). (**M**, **N**) Scatter plots documenting the lack of correlation between the basal ratios and EC_50_ of individual ETI-responsive glomeruli in WT and AD mice of different ages (Pearson correlation coefficients: 0.06-0.22, P=0.27-0.77, n=17-37 glomeruli).

**Fig. 6.**
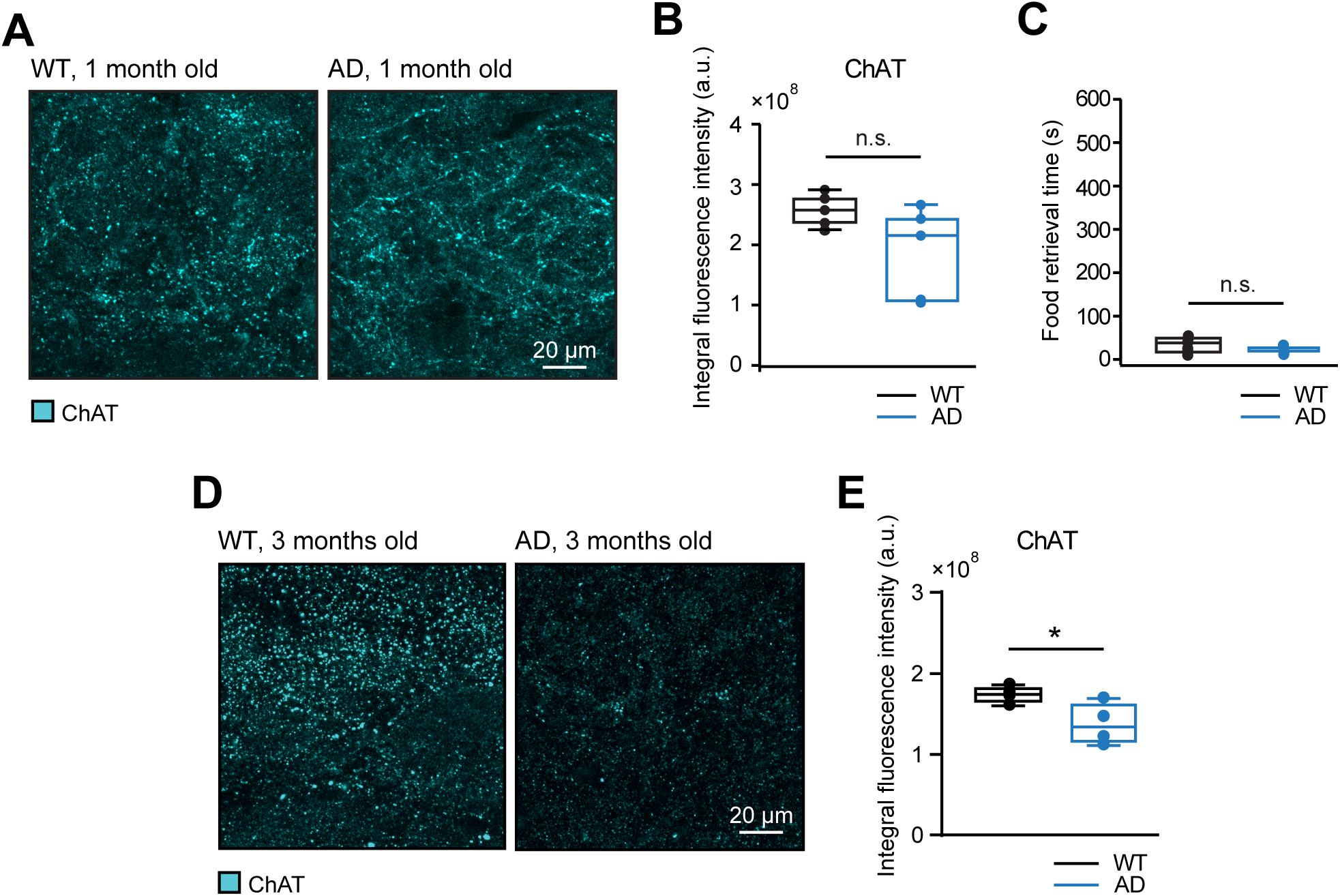
Gradual reduction of cholinergic innervation of the glomerular layer in a mouse model for Alzheimer’s disease. (**A**) Sample maximum intensity projection images (1-12 µm depth, 1 µm step) of the glomerular layer in fixed OB slices from 1-month-old WT and AD mice, labeled with antibodies against choline acetyltransferase (ChAT). (**B)** Box plots showing mean (per mouse) background-subtracted integral ChAT fluorescence intensity values (see Materials and methods) of different fields of view (FOVs) in 1-month-old WT and AD mice (Wilcoxon Rank test, P=0.12, n=5 WT and 5 AD mice, 5 FOVs/mouse). (**C**) Box plots illustrating the time spent to retrieve the chocolate in a buried food pellet test by 1-month-old WT and AD mice (Wilcoxon Rank test, P=0.3, n=7 WT and 5 AD mice). (**D**, **E**) The same analyses as in (**A**, **B**) for 3-4 months old mice (**D**: 0-11 µm depth, 1 µm step; **E**: t-test, P=0.04, n=4 WT (22 FOVs) and 4 AD (21 FOVs) mice).

The observed above differences were much less pronounced in the output (mitral) cell layer of the OB. Here, we analyzed a single age group (4-7 months of age), as the data obtained in the glomerular layer were similar in 3-4 and 5-7 months old mice. The *in vivo* basal ratios of mitral cells in WT (n=74 cells, 4 mice) and AD (n=90 cells, 4 mice) animals were only slightly (although significantly) different (fig. S7 A to C), with higher values observed in mitral/tufted cells of AD mice in line with less inhibitory drive from the glomerular cell layer (Fig. 5). Of note, the intercellular variance in basal ratios did not differ significantly between the 2 genotypes (fig. S7D). Interestingly, in AD mice, the density of cholinergic projections to the glomerular layer of the bulb, visualized immunohistochemically with an anti-ChAT (choline acetyltransferase) antibody, was similar to that in WT mice at 1 month of age but decreased significantly already at 3-4 months of age (Fig. 6, A to B and D to E). Consistently, the 1-month-old AD and WT mice performed similarly well in a buried food pellet test (Fig. 6C), with AD-related deficits becoming obvious from 3 months onwards (Fig. 5, E and K).

Thus, the early impairment of olfaction in 3-4-month-old AD mice is likely caused by the reduced inhomogeneity of the glomerular map due to the decrease in cholinergic innervation. As the disease develops, decreased odor-sensitivity of the olfactory system further worsens the animal’s ability to detect odors.

## Discussion

Here, we directly visualize the glomerular activity map in awake head-restrained mice and show its striking inter-glomerular inhomogeneity. Within a given glomerulus, however, the activity levels were rather similar, underscoring the “functional unit” properties of the glomeruli, previously seen when analyzing their odor-evoked responses (*18, 35*). Under resting awake conditions, the glomerular activity maps were impressively stable over days and weeks, but changed considerably under different types of anesthesia and blockage of cholinergic centrifugal inputs, indicating that they are brain state-dependent. According to our data, the glomerulus-specific activity levels were sustained by the combination of (i) ongoing endogenous spiking of olfactory sensory neurons, (ii) air flow-stimulated OSN firing, (iii) modulatory cholinergic projections to the olfactory bulb and, to a small extent, (iv) by the ambient odor environment. Importantly, none of the aforementioned mechanisms alone was able to sustain the observed levels of inhomogeneity. In a mouse model of AD, loss of glomerular map inhomogeneity developed very early, in parallel with a decrease in centrifugal cholinergic projections and before the amyloid plaque deposition in the OB. Moreover, it was paralleled by an impairment in olfactory-driven behavior, thereby identifying the glomerular map inhomogeneity as a novel AD biomarker.

The described molecular mechanisms underlying the glomerular map inhomogeneity align well with previous data obtained in various types of preparations. Dissociated OSNs, OSNs in the isolated olfactory epithelium or olfactory epithelium slices, for example, are known to be endogenously active with spiking frequencies homogeneously distributed between 0 and 10 Hz (*12–14, 16, 36*). Moreover, cells expressing different types of odorant receptors are broadly tuned to different spiking frequencies and show different degrees of mechanosensitivity (*13, 16, 36, 37*).

When stimulated by the patterned (artificial sniffing) or un-patterned airflow in anesthetized/awake head-fixed rodents, OSN axons showed airflow-induced responses in ∼ 50% of all glomeruli (*38, 39*). The amplitudes of these airflow-driven responses varied between individual glomeruli, suggesting that the extent of mechanosensitivity is olfactory receptor-specific. Interestingly, in the same preparation (*39*), the dendrites of airflow-responsive mitral cells projected to 80% of all glomeruli, hinting towards the mechanisms enhancing airflow-induced responsiveness beyond the input layer of the OB. Besides, the OSN inputs can be further modified by the intrinsically bursting external tufted cells, entrained by these inputs and synapsing on excitatory mitral cells, external tufted cells, and superficial tufted cells, as well as inhibitory periglomerular neurons (*40, 41*).

The literature concerning the cholinergic modulation in the OB, often stemming from electrophysiological recording in anesthetized preparations, remains so far inconclusive. Thus, electrical or optogenetic stimulation of the cholinergic neurons in the horizontal limb of the diagonal band of Broca (HDB), the main source of the cholinergic projections to the OB (*42, 43*), caused inhibition (*44*) or excitation (*42, 45*) of the output mitral/tufted cells as well as inhibition of glomerular and granule cell layer interneurons (*44*). Besides, the stimulation also modulated odor responses of individual glomeruli in different directions (*42*). Moreover, opposite effects were exerted by different subtypes of acetylcholine receptors (out of 7 subtypes of nicotinic and 2 types of muscarinic acetylcholine receptors detected in the OB), with nicotinic receptors activating mitral, periglomerular and external tufted cells, and muscarinic receptors mediating either granule cells excitation (M1 receptors), increasing GABA release from granule onto mitral cells, or granule cells inhibition (M2 receptors) (*42, 43, 46*). However, when optogenetically exciting only cholinergic axons projecting to the OB and not all cholinergic neurons in the HDB, the spontaneous firing of mitral cells as well as their responses to artificial inhalation of clean air were enhanced (*47*), well in line with a strong reduction of the glomerular map activity observed under cholinergic receptor blockers in our study (Fig. 4E-H). In fact, the latter treatment completely silenced 50% of all glomeruli, consistent with the dense cholinergic innervation of both glomerular and mitral cell layers of rodents and primates (Fig.6; (*45, 47, 48*)).

Our data unify all the scattered information described above by introducing and directly visualizing the extremely stable inhomogeneous map of the basal glomerular activity as a background against which olfactory signal processing is performed. The functional significance of this inhomogeneous background was assessed in a mouse model of AD, in which the loss of basal map inhomogeneity occurred before the amyloid plaque deposition and the reduction in the glomerular odor sensitivity. Remarkably, it was paralleled by the early loss of cholinergic innervation of the glomerular layer and was associated with the impairment of odor detection ability, thus providing a mechanistic explanation for early olfactory impairment in AD patients. The young age at which the loss of cholinergic innervation and odor detection ability jointly occur (between 1 and 3-4 months of age) makes other potential explanations (e.g., neurodegeneration in odor-sensing or processing pathways) unlikely. Of note, mouse models of AD expressing mutant APP and presenilin proteins do not recapitulate neurodegeneration seen in humans (*49*). The observed association between basal glomerular activity and odor sensing is in line with the poor odor discrimination ability of mice with reduced ongoing activity due to overexpression of Kir2.1 channels in OSNs (*12*). Thus, the resting state glomerular activity map adds another layer to the complex computational network of the OB, increasing the dynamic range of and the variance in the system via glomerulus-specific functional inhomogeneity. Moreover, it enables responding to odorants not only with graded increases (ON-glomeruli) but also with graded decreases (Inhibited-glomeruli, (*18*)) in glomerular activity. The importance of this new layer for signal processing is emphasized by computational/machine learning studies suggesting that functional inhomogeneity in neural networks allows for an increase in computational power and performance at low cost (metabolic efficiency), promotes and stabilizes robust learning (computational efficiency), improves generalization and enhances pattern separation (*50–52*).

## Materials and methods

### Ethics statement

In this study, we used C57BL/6N wildtype (WT) mice and APPswe/PS1_G384A_ mice of either sex, bred in our animal facility. Neurons of APPswe/PS1_G384A_ mice express amyloid precursor protein (APP) and presenilin 1 (PS1) with mutations (*32*), known to cause Alzheimer’s disease in humans. Female mice were housed in groups of 3-5, while male mice were conventionally housed in individual neighboring cages with all olfactory, visual, and acoustic stimuli preserved, to avoid fights and injuries. Unless otherwise indicated, all animals were kept under a 12-hour light/dark cycle with *ad libitum* access to food and water. All experiments were conducted following institutional animal welfare guidelines and were approved by the state government of Baden-Wuerttemberg, Germany.

### Labeling of the olfactory bulb with the Ca^2+^ indicator Twitch-2B

To label neural networks in the olfactory bulb, an AAV-based viral vector encoding the FRET-based Ca^2+^ indicator Twitch-2B (Kd = 200 nM) under the human Synapsin1 promoter (AAV1-hSyn1-Twitch-2B, Addgene viral prep # 100040-AAV1) was used (*53*). The virus was diluted 1:7, and 1µl of the virus-containing solution was stereotactically injected mediocaudally (at 0.35 mm, 0.25 mm, and 0.15 mm depth (∼300 nl per injection) and a 45° angle from the horizontal plane) into both olfactory hemibulbs of 2-to-6-month-old mice, followed by chronic cranial window implantation (see below). This enabled labeling of both pre- (fig. S8) and postsynaptic (fig. S7, (*19*)) glomerular structures with Twitch-2B.

Compared to single-wavelength Ca^2+^ indicators like GCaMPs, the use of ratiometric FRET-based indicators, including Twitch-2B, provides more accurate measurements better suited for long-term *in vivo* functional imaging studies, because they are less influenced by changes in the optical path length, excitation light intensity, indicator expression level, as well as by movement artefacts. Moreover, Twitch-2B is substantially brighter than GCaMPs in non-spiking neurons, allowing better identification of expressing cells and their subcellular structures, has a better linearity as well as reduced Ca^2+^ buffering capacity and cytotoxicity, due to a reduced number of Ca^2+^ binding sites (2 in Twitch-2B vs. 4 in GCaMPs; (*53*)).

### Chronic cranial window implantation

Surgery was performed as described earlier (*19, 54*). Briefly, mice were anesthetized with an intraperitoneal (i.p.) injection of midazolam (5 mg/kg BW), medetomidine (0.5 mg/kg BW), and fentanyl (0.05 mg/kg BW), hereafter abbreviated as ‘MMF’ anesthesia. The surgical depth of the anesthesia was confirmed by the absence of the toe pinch reflex. Before surgery, an eye ointment (Bepanthen, Bayer, Leverkusen) was applied to prevent drying out of the cornea. Besides, xylocaine (2% w/v) was injected subcutaneously (s.c.) to induce local anesthesia at the incision site, and dexamethasone (0.2 mg/kg BW, Sigma-Aldrich Germany) was injected s.c. to prevent swelling of the brain upon removal of the bone. The body temperature was monitored using a rectal temperature probe and maintained between 36-37°C. After removing the skin above the OB, two craniotomies were made for each olfactory hemibulb using a dental drill (NSK, Ultimate 500): the region around both OB hemispheres and the midline bone covering the olfactory sinus between the hemispheres was thinned until two loosely attached islands formed. Subsequently, ringer injection solution (B. Braun, Melsungen AG) was applied to prevent drying out of the brain surface, and the islands were gently removed with forceps while the midline bone above the olfactory sinus, as well as the dura were left intact. Both hemibulbs were covered with one round coverslip (Ø 3 mm, Warner Instruments, Hamden, CT). The remaining exposed skull was covered with dental cement (Tetric EvoFlow, Ivoclar Vivadent AG), and a small stainless steel or titanium holder was fixed to the skull caudally to the coverslip. Postoperative care included an analgesic dose of carprofen (5 mg/kg BW, Pfizer GmbH) for 3 days s.c. and 0.025% of the antibiotic enrofloxacin (Baytril, Bayer) for 10 days in drinking water. The two-photon imaging commenced 3-4 weeks after the surgery. Before the first awake two-photon imaging session, mice were habituated to fixation in the imaging setup for 10-15 days by daily fixations, lasting between a few seconds (at the beginning of training) and up to 90 minutes (at the end of training).

For applying receptor antagonists to the surface of the olfactory bulb, trained mice were implanted with the chronic cranial window as detailed above, but the coverslip contained a 1.0×0.1 mm slit (*20*), located rostrally above the cavity between the OB hemispheres. At the end of the surgery, the slit was covered with Kwik-Cast and sealed on top with Kwik-Sil (both are silicon elastomers from World Precision Instruments Germany GmbH, Berlin), and imaging experiments commenced one day later. The silicon elastomers were removed before the drug application.

### In vivo two-photon calcium imaging

#### Monitoring of mouse physiological parameters and anesthesia induction

Only trained mice, habituated to the setup and the head fixation, were used. In awake mice, the respiration rate was monitored using a thermistor (Murata NCP15XH103J03RC; Conrad) positioned in front of one nostril. In anesthetized mice, the respiration rate was monitored with a pressure sensor (ADInstruments, Dunedin, New Zealand) attached to the back of the mouse. Under isoflurane anesthesia, isoflurane concentration was maintained at 0.9-1.5% in O_2_ to keep the breathing rates between 110 and 160 breaths per minute. Otherwise, mice were injected i.p. with ketamine/xylazine (80/4 µg per g BW) or MMF (composition see above). Animals breathed freely throughout the experiments. Two-photon imaging started 10-15 min after anesthesia induction.

#### Imaging setup and image acquisition

We used an Olympus Fluoview 1000 laser scanning microscope (Olympus Europa GmbH) coupled to a mode-locked Ti:Sapphire laser (Mai Tai Deep See, Spectra Physics GmbH) and operating at 690-1040 nm with a pulse width of <100 fs and a repetition rate of 80 MHz. Images were acquired using an excitation wavelength of 890 nm and a 20x Plan-Apochromat 1.0 NA water-immersion objective (Carl Zeiss AG). Emitted light was split using a 515 nm dichroic mirror and a 475/64 nm bandpass or a 500 nm long pass filter for mCerulean3 and cpVenus^CD^ channels, respectively. For time series, three consecutive sequences of 80 (Fig. 5) or 1000 (fig. S3) frames at a rate of 4-7.8 Hz with an inter-sequence interval of 30 to 60 seconds, and an image size of 512×256 pixels were recorded at a single depth level within the glomerular layer. For the 3-dimensional reconstruction of the glomerular layer, images were taken from the dura mater to a depth of around 120 μm with an image size of 640×640 pixels, using a Kalman filter of 3 and a step size of 2 µm.

Immunohistochemically labeled OB slices were imaged through a 40x water immersion objective (0.80 NA, Nikon, Tokyo, Japan) at 800 nm excitation wavelength. A 585 nm dichroic mirror was used to split the light emitted by the two Alexa dyes. A 536/40 nm band-pass and a 585 nm long-pass filter were used to filter the Alexa Fluor 488 and the Alexa Fluor 594 emission light, respectively. For capturing the Thioflavin-S fluorescence, a 515 nm dichroic mirror and a 512 nm short-pass filter were used.

### Odorant delivery

In the majority of experiments, glomerular basal Twitch-2B ratios were measured in mice freely breathing the ambient air of a ventilated room, termed “Room air”. In some experiments, the room air was replaced by “Clean air” from an air tank, by placing a custom-built flow-dilution olfactometer (similar design as in ref. (*18*)) approximately one centimeter in front of the mouse’s snout and providing a constant clean airflow at the rate of 300 ml/min. To record odor-evoked Ca^2+^ signals, ethyl tiglate (ETI), ethyl butyrate (EBU), or isoamyl acetate (IAA), all known to activate the dorsal OB (*55*), were applied via the flow-dilution olfactometer, in which the pure air was mixed with the saturated odorant vapor. For measuring the dose-response curves (Fig. 5), the following odorant concentrations (% of s.v.) were used: 0.07, 0.11, 0.2, 0.34, 0.68, 1.01, 1.7, 2.64, 4.86, and 7.89. For each concentration, the odorants were delivered for 4 seconds in three trials with an inter-trial interval of 1-2 minutes. The resulting dose-response curves were used to extract the effective odor concentrations that elicit half-maximal responses (EC_50_). For continuous odor application over 30 minutes, the odorant concentration of 0.18% s.v. (“Clear air + long odor on”) was used. All odorants were purchased from Sigma-Aldrich Germany in the highest commercially available purity.

### Local application of antagonists in awake mice

As inflammation-induced changes of the tissue function were reported to begin 2 days after surgery (*56, 57*) and because the dura mater in our preparations became impermeable to drugs after 4 weeks (own observation), we applied receptor antagonists 12-24 hours after surgery. Drugs (all from Sigma-Aldrich) were diluted in HEPES-buffered Ringer solution (in mM: 150 NaCl, 4.5 KCl, 10 HEPES, 1MgCl_2_, 1.6 CaCl_2_, pH 7.4). A drop of around 40 µl was applied onto the cover glass with a slit. The following drugs/concentrations were used: the voltage-gated Na^+^ channel blocker tetrodotoxin (TTX, 5 µM), the nonselective α1- and α2- adrenergic receptor antagonist prazosin (100 µM), the nonselective 5-HT_1_-, 5-HT_2_-, 5-HT_7_- serotonin receptor antagonist methysergide (4 mM). The mAChR-antagonist scopolamine and the nAChR-antagonist mecamylamine were prepared first in HEPES-buffered Ringer solution and then mixed to a final concentration of 50 mM and 115 mM, respectively. To ensure that sufficient drug concentrations reached the target cells via diffusion through the small slit, we used higher concentrations than the ones described previously in anesthetized mice, when perfusing a wide area of the OB surface (*24, 47*). The initial imaging session was performed with the Kwik-Cast/Sil still covering the slit (see ref. (*19*) for further details). Then, Kwik-Cast/Sil was gently removed to apply the HEPES-buffered Ringer solution (Fig. 4, Control) and subsequently, the receptor blockers (Fig. 4, Block).

### Modulating the activity of olfactory sensory neurons

To address the contribution of olfactory sensory neurons (OSN) to the overall basal glomerular activity levels, two-photon imaging experiments were performed in awake animals. Only trained mice, habituated to the setup and the head fixation routine, were used. After recording basal glomerular Twitch-2B ratios (Control) and odor-induced responses (fig. S4, details see below), mice were returned to their home cage for five minutes. Thereafter, the OSNs were targeted in one of three different ways, as described below.

#### Non-invasive intranasal application of TTX

To block OSN activity within the sensory epithelium, a non-invasive intranasal application of TTX (50 µM in PBS) was used. 10 µl of the TTX-containing solution was applied twice as droplets directly into one nostril of mice, which were briefly (≤ 2 minutes) anesthetized with isoflurane. Mice were returned to their home cages and allowed to rest for 10 minutes to recover from anesthesia. Thereafter, a second identical set of TTX applications was performed in the awake state. Immediately after the application of the blocker, the mice were head-fixed in the setup, and the same region of interest as under control conditions was imaged.

#### Dichlobenil administration

For chemical ablation of olfactory sensory neurons, the known olfactotoxic substance dichlobenil (**2,6-Dichlorobenzonitrile** (*21*)) was administered via a single i.p. injection (50 µg/g body weight in 1 µl/g body weight DMSO) after the control imaging session. The effect of dichlobenil was monitored over time, with imaging sessions at 3, 6, 9, 12, 24, and 48 h after dichlobenil administration. Between the imaging sessions, mice were transferred to their home cage.

#### Nose plugging

For blocking the activity of OSNs caused by the air flow-induced mechanosensation and ambient odors, one nostril was reversibly plugged by the silicon elastomer Kwik-Cast under short (5-8 minutes) isoflurane anesthesia. To do so, the mouse was placed on its back on a heating pad, and the two-component liquid elastomer was carefully applied to the nostril. The anesthesia was maintained, and the viscose elastomer was allowed to harden, thus forming a tight-fitting plug. Immediately thereafter, the isoflurane supply was interrupted, and the mouse was transferred to the setup to be imaged at different time points (Fig. 3). We used a plastic collar to prevent the awake animal from removing the plug. After the measurement, the collar was removed, allowing the mice to remove the plug within 1-2 minutes. After reopening the nostril, the imaging session resumed.

### Buried food pellet test

To test the ability of mice, exposed overnight to 16 hours of food deprivation, to detect odors, we measured the time mice spent retrieving 50 mg of chocolate, hidden under 1 cm of bedding material (*34*). Mice were habituated to the new clean cage for 5 min and then returned for 30 min to their home cage before the test. In the meantime, the chocolate was randomly placed in one of the four cage corners. Subsequently, the mouse was placed in the center of the cage, and the time needed to retrieve the chocolate (retrieval time) by grasping it with front paws or teeth was recorded. The single test trial lasted 8 min. A failure was recorded if the animal failed to retrieve the chocolate within this time.

### Immunohistochemistry

The fixed OB tissue of 1, 3-4 or 5-7 months old APPswe/PS1_G384A_ mice was taken from our institute’s biobank, cut with a cryostat (Leica, Germany) into 50-μm-thick sagittal sections, and washed 3×10 min with PBS. The free-floating OB slices were blocked with 1% BSA (Bovine Serum Albumin, Fraktion V, Serva), 10% normal donkey serum (Dianova, 017-000-121), and 1% Triton X-100 (Sigma, T-9284) in PBS for 2 h at room temperature. Next, the slices were incubated at 4 °C for 48 hours with anti-choline acetyltransferase antibodies (ChAT, 1:250, Novus, NBP1-30052) diluted in another blocking solution containing 0.5 % BSA, 10% normal donkey serum, and 1 % Triton X-100 in PBS. Subsequently, the slices were washed 5×10 min in PBS and then incubated for 2 h in the darkness in PBS containing 2 % BSA and Alexa-594 conjugated secondary antibodies (1:1000) at room temperature. Then, the slices were washed 5×10 min in PBS in the darkness and mounted in Vectashield Mounting Medium (Vector Laboratories, Burlingame, CA). To obtain the data shown in the fig. S6 or S8, the free-floating OB slices were treated with a blocking solution (5% normal donkey serum and 1% Triton-X in PBS) for 1 h before overnight incubation at 4 °C with either the primary anti-Iba-1 (1:500, Wako, USA, 019-19741) and anti-CD68 (1:1000, Abd Serotec, UK, MCA1957) or the primary goat-anti-OMP (Olfactory marker protein; 1:1000, Wako, USA, 544-10001) and rabbit-anti-GFP (1:1500, Rockland, USA, 600-401-215) antibodies, diluted in blocking solution. After rinsing the slices 3×10 min in PBS, slices were incubated in the darkness in a 2% BSA solution containing Alexa-488 and Alexa-594 conjugated secondary antibodies (1:1000 for both Alexa-488 and Alexa-594) for 2 h at room temperature. Thereafter, the slices used for fig. S6 were transferred for 8 minutes into a solution containing 2*10^-6^ % Thioflavin S in PBS. Finally, all slices were washed 3×10 min in PBS in the darkness and mounted on fluorescence-free Superfrost microscope glass slides in Vectashield mounting medium.

### Data analysis

#### Time series of basal and odor-evoked Ca^2+^ signals

For analysis of glomerular signals, regions of interest (ROIs) were drawn manually in Fiji (https://imagej.net/Fiji). A reference background ROI was drawn in the darkest spot of the image. Further analyses were performed using custom-written scripts in MATLAB (The MathWorks, Inc.). The respective mean background intensity was individually subtracted from the mean fluorescence intensities in mCerulean3 and cpVenus^CD^ channels before cpVenus^CD^/mCerulean3 ratios were calculated for all ROIs. Odor-evoked Ca^2+^ transients were counted as responses when their ΔR/R signal was six times larger than the standard deviation of the corresponding baseline noise, and when a minimum of 15% ΔR/R was reached.

#### Three-dimensional measurements of basal glomerular Twitch-2B ratios

We used a custom-written MATLAB script to load 3D stacks, draw, for each glomerulus, eight ROIs at different levels covering the entire glomerular volume, and extract mean fluorescence intensity values of mCerulean3 and cpVenus^CD^. The background fluorescence was subtracted individually for each channel and z-level, and a ratio of cpVenus^CD^/mCerulean3 was calculated. The resulting median ratio values of the eight z-levels were taken as the basal ratio value of an individual glomerulus.

#### Computation of color-coded ratio maps (color maps)

To compute the color-coded ratio maps, each full-frame image (640×640 pixels) of the gray-scale 3D image stack was filtered with a 7×7 average filter. Then the data was analyzed similarly to the procedure described above, but the ratios were calculated per pixel for those pixels in the image whose mean intensities were between 100 and 2500. The remaining pixels were blanked to avoid over- (when dividing by a small number) or underestimation (when including the saturated pixels of cell somata) of ratio values. The resulting ratio images were color-coded with a colorblind-friendly continuous color map (turbo; (https://research.google/blog/turbo-an-improved-rainbow-colormap-for-visualization/)).

#### Calculation of the effect size

In experiments shown in Figs. 2 to 4, a reduction of the basal Twitch-2B ratio was calculated as a percentage of the maximal block that can be reached, termed “effect size”. The maximum theoretically possible block was defined as the reduction towards the lowest glomerular ratio level (1.3) ever measured under the TTX application to the surface of the olfactory bulb (fig. S2). We calculated the difference between the basal ratio measured in the control condition and 1.3 (R_ctr_ - 1.3) and the difference between the basal ratios measured in the control and the test condition (R_ctr_ - R_test_). The effect size (in %) was then calculated according to the formula:

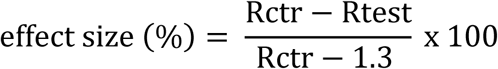

#### Analyses of the integral ChAT fluorescence intensity

Maximum intensity projection (MIP) images (3D stacks: 122 x 122 µm, 10 µm depth, 1 μm step) of the glomerular cell layer, taken under identical imaging conditions, were imported into Igor (WaveMetrics, Portland, US) to calculate the cumulative intensity histograms. The intensity of the background fluorescence was estimated based on 3 MIP images, taken in corresponding control slices (labeled by secondary antibody only). To do so, we averaged the readouts of three 90^th^ percentiles of the control cumulative intensity histograms and used this value for background subtraction to obtain the background-subtracted integral ChAT fluorescence intensity for each field of view (FOV).

#### Analyses of heterogeneity and colocalized tissue volume

The inter-glomerular variance of Twitch-2B ratios, as a measure of heterogeneity, was calculated for each image stack, as follows:

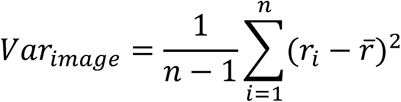

where *r_i_* is the Twitch-2B ratio of the 𝑖^𝑡ℎ^ glomerulus in the given image stack, and *r̄* is the mean of Twitch-2B ratios across all glomeruli (𝑛) in the stack. We used the exact two-sided weighted Mann-Whitney-Wilcoxon test (*58*) to test for inhomogeneity differences between AD and WT mice in two age groups. The number of glomeruli per image stack was used as a weight to adjust for the varying numbers of glomeruli across different stacks.

The inter-cellular variance of mitral/tufted cells was also calculated using the above equation, where *r_i_* denotes Twitch-2B ratio of the 𝑖^𝑡ℎ^ cell and *r̄* is the mean Twitch-2B ratio across all *n* cells. Inhomogeneity differences between AD and WT mice were tested with the exact two-sided Mann-Whitney-Wilcoxon test.

For analyses of data shown in the fig. S8, surface rendering algorithm was used to mask both Twitch-2B (anti-GFP labeling) and OMP channels in Imaris (version 10.2.0). The resulting surfaces were automatically thresholded with the ImarisColoc module, and the fraction of the overlapping volume was calculated as below:

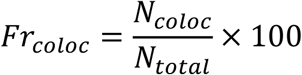

where *Fr_coloc_* is the percentage of the overlapping volume, *N_coloc_* is the number of the colocalized voxels, and *N_total_* denotes the total number of voxels in the Twitch-2B channel.

#### Statistical analysis and data presentation

Statistical tests were performed in MATLAB, GraphPad Prism (GraphPad Software, www.graphpad.com), Igor (WaveMetrics, Portland, US), R version 4.2.1, or Vassar Stats (website for statistical computation, http://vassarstats.net/). The one-sample Kolmogorov-Smirnov test was used to check for the normality of data distribution. The Levene test was used to check for homoscedasticity. Non-parametric tests were performed if parametric test assumptions were violated. In cases with a repeated measure design and ANOVA assumption violations (Figs. 1F, 2C, 2F, and 3D), we performed the Friedman test followed by the Conover post-hoc test and Bonferroni correction, considering the control values as a separate group. All statistical tests were two-sided. The P values smaller than 0.05 were considered significant. Unless otherwise indicated, data are presented as median ± interquartile range. Lines of boxes in box plots represent 25^th^ and 75^th^ percentiles, and whiskers show the minimum and maximum data points, respectively. All bar, box, cumulative, or x-y scatter plots were calculated and plotted with custom-written MATLAB scripts and displayed with Adobe Illustrator CC 2017.

## Acknowledgments

We thank E. Zirdum, A. Weible, K. Schönntag, B. Ott and K. Schmidt for technical assistance; Saib Rinnawi for some data shown in fig. S4 and Nima Mojtahedi, Marie-Estelle Schmidt, Pariya Khodabakhsh and Joe Sheppard for help with Matlab coding, Imaris-assisted analyses, and data handling.

## Author contributions

Conceptualization: O.G.

Investigation: S.F., N.F-T, W-Y.T., Y.K.

Methodology: Y.K., N.F-T., S.F.

Supervision: O.G.

Visualization: S.F., N.F-T., W-Y.T., O.G.

Statistical analyses: S.F., N.F-T, W-Y.T., Y.K., F.K.

Writing – original draft: S.F., N.F-T., O.G.

Writing – review and editing: O.G., S.F., N.F-T., W-Y.T., F.K., Y.K.

## Data and materials availability

All data are available in the main text or the supplementary materials.

