## Supplemental Data for "Glomerulus-specific inhomogeneity of the basal activity map in the olfactory bulb supports olfactory-driven behavior"

Stefan Fink *et al.*

**This PDF file includes:**

Figs. S1 to S8

#### Supplementary Figures

**Fig. S1. Stability of glomerular basal Twitch-2B ratios over longer time periods.** Color maps illustrating the basal Twitch-2B ratios of different glomeruli measured in the awake state on different experimental days (as indicated in the image).

**Fig. S2. Defining glomerular activity levels.** (**A**) Color maps, illustrating the level of basal glomerular activity before (Control) and after application of 5 μM tetrodotoxin (TTX) to the surface of the olfactory bulb through the cranial window with a slit, located in the lower left corner. (**B**) Histogram, illustrating the distribution of basal ratios of individual glomeruli before (white bars) and during (blue bars) TTX application (n=22 glomeruli, 3 mice). (**C**) Probability density functions of the distributions, shown in (B). Note that under TTX all values in (B) and (C) lay below 1.8 (broken line in (C)). Therefore, all glomeruli with basal ratios below 1.8 were considered non-active (comprising non-spiking cells). (**D**) Box plot showing the distribution of basal Twitch-2B ratios before and during TTX application (paired t-test, $P=1.66*10^{-8}$, n=22 glomeruli, 3 mice).

**Fig. S3 Glomerular basal activity map in awake mice imaged at higher temporal resolution.** **(A)** A sample color map acquired at 0.22 Hz sampling rate. **(B)** Average intensity projection of a time series recorded at 7.76 Hz from the area marked with a square in (A). **(C)** $Ca^{2+}$-traces recorded from the glomeruli (1-4) delineated in (A) and (B). **(D, E)** To compare the two approaches, the time series were binned (down-sampled) to the same sampling rate as recorded in (A) by averaging 36 image frames for each epoch (broken lines in (E)). The resulting mean values (red squares) were taken as the glomerular basal activity level in each epoch. **(D)** The sample color map computed from a single epoch marked in (E). **(F)** Summary of similar comparisons as in (A-E) conducted for 19 glomeruli from 4 mice. For each slow (0.22 Hz) or fast (7.76 Hz) measurement, median ratio values were taken.

**Fig. S4. Intranasal TTX application inhibits odor-evoked glomerular responses in the ipsilateral hemibulb.** (**A**) Left panel: average intensity projection of glomeruli responding to ethyl butyrate (EBU, 1.7% s.v.). Right panel: overlay of traces illustrating odor-evoked $Ca^{2+}$-transients in individual glomeruli (marked with a respective number on the left image) before (Control), during (TTX), and 24 hours after (Recovery) the non-invasive intranasal TTX application. (**B**) Odor-responsive glomeruli were identified using the Frame Subtraction procedure (NeuroPlex software, RedShirt Imaging, Decatur, GA, USA). Frame subtraction images (obtained by subtracting the average of 15 frames recorded in the mCerulean3 channel during the odorant presentation from the average of 15 frames recorded before the odorant presentation) are shown for control, TTX, and recovery conditions. The scale is shown in arbitrary units (AU). Brighter areas represent odor-responsive cells and glomeruli.

**Fig. S5. Small but significant absolute ratio changes caused by the long odorant application.** Box plot showing the median absolute ratio changes measured during the 30-min-long period in Room air, Clean air, and Clean air + Odor (n=6, 4, 6 mice, respectively). One-way ANOVA $P=4.8*10^{-5}$, F=23.47, post-hoc Tukey HSD test (Room vs. Clean air: P=0.19, Clean air vs. Clean air + Odor: $P=2.9*10^{-3}$, Room air vs. Clean air + Odor: $P=3.9*10^{-5}$).

**Fig. S6. Amyloid β deposition in the olfactory bulb of APPswe/PS1G384A mice.** Sample MIP images, obtained from fixed OB slices of 3 (top) and 5 (bottom) months old AD mice and stained against a microglia/macrophage marker Iba-1 (green) and an endosomal/lysosomal marker CD68 (red). Amyloid plaques, labeled by Thioflavin S, are shown in blue. Merged images are shown in the right column. Note that plaques (arrowheads) are seen only in 5-month-old mice and are restricted to the granule cell layer.

**Fig. S7. Little difference in the basal Twitch-2B ratios of mitral/tufted cells and their intercellular variance between WT and AD mice.** (**A, B**) Sample average intensity projection images (5 µm depth, step 1 µm, > 200 µm below the surface) of the dorsal OB labeled with the $Ca^{2+}$-indicator Twitch-2B (left) and the corresponding color maps (right) taken in 4-7-months old WT and AD mice. Arrowheads point to mitral/tufted cells, identified by their size (≥ 15 µm in diameter) and their location below the external plexiform layer. (**C**) Cumulative probability plots illustrating distributions of basal Twitch-2B ratios in WT and AD mice of respective ages (Kolmogorov-Smirnov test, P=0.02, n=74 cells from 4 WT and 90 cells from 4 AD mice). (**D**) Box plots illustrating the intercellular variances (calculated as described in Materials and methods) in WT and AD mice (exact weighted Mann-Whitney-Wilcoxon test, P=0.69, n=4 WT and 4 AD mice).

**Fig. S8. Presynaptic structures contribute to the glomerular Twitch-2B signal.** (**A**) Sample MIP images, obtained from fixed OB slices of a WT mouse and stained against GFP (green, visualizing Twitch-2B) and a marker of the olfactory sensory neurons, Olfactory marker protein (OMP, yellow). The right panel shows all overlapping voxels in a 35-µm-thick volume

(Imaris-assisted analysis described in Materials and methods). (B) Box plots illustrating fractions of voxels with overlapping Twitch-2B and OMP staining among all OMP-positive voxels (n=7 mice).

### Fig. S1. Stability of glomerular basal Twitch-2B ratios over longer time periods

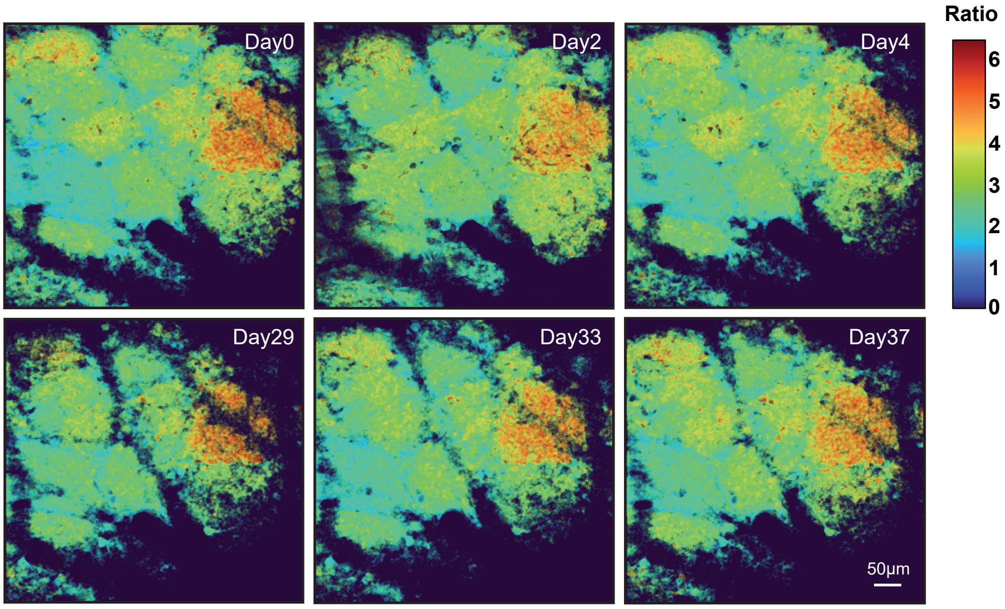

#### Fig. S2. Defining glomerular activity levels

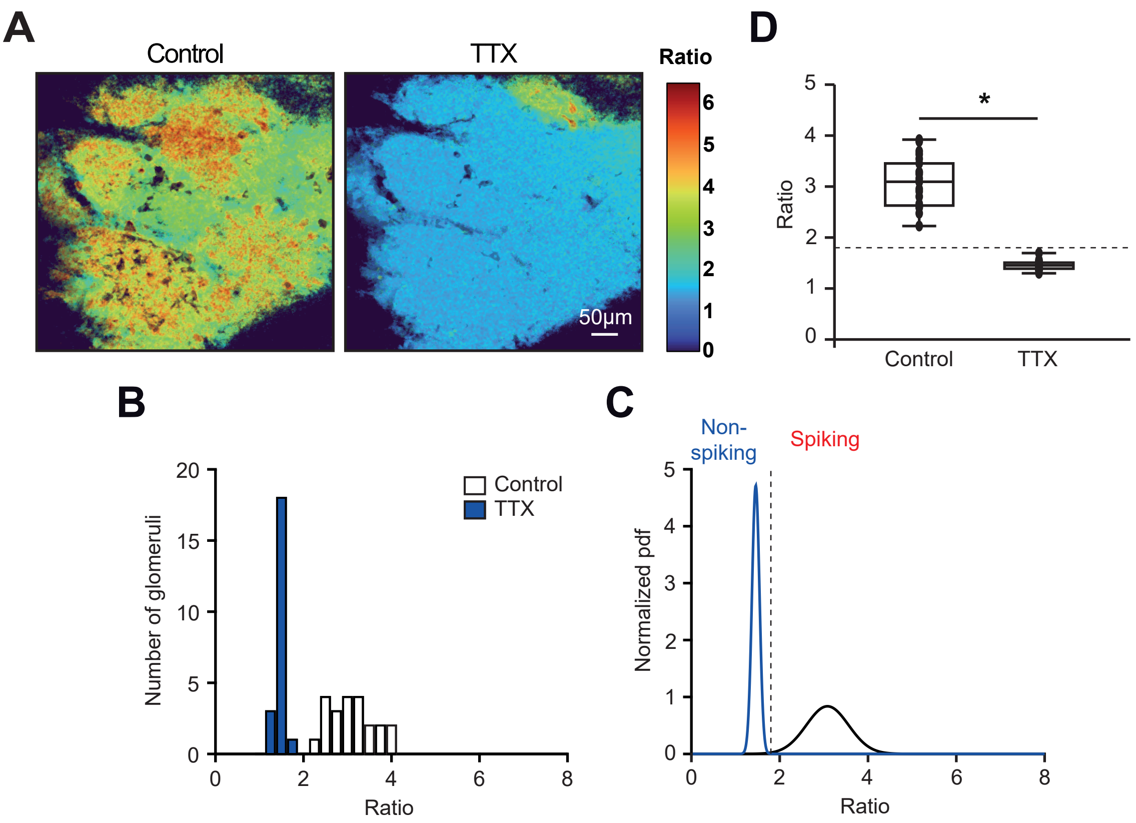

**Fig. S3. Glomerular activity maps imaged at higher temporal resolution**

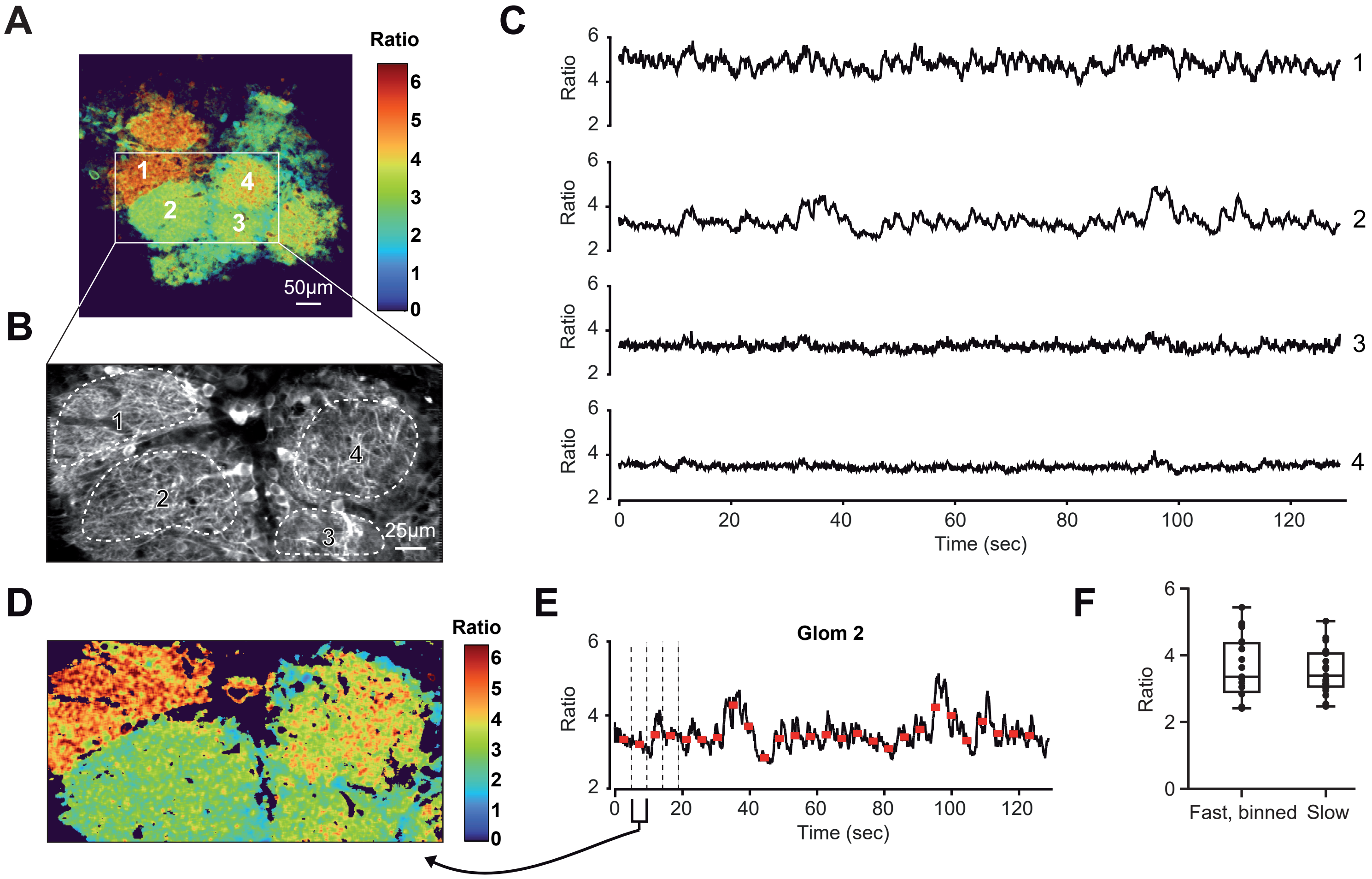

**Fig. S4. Intranasal TTX application inhibits odor-evoked glomerular responses in the ipsilateral hemibulb**

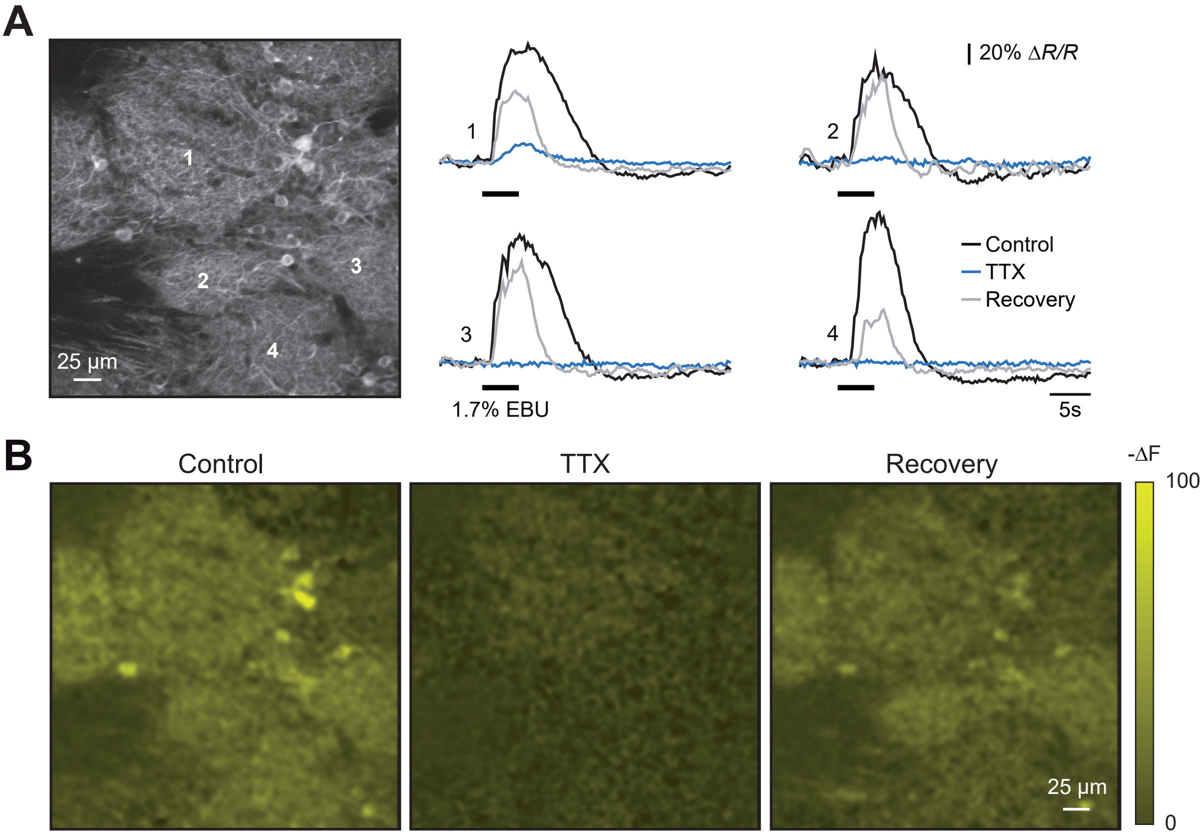

**Fig. S5. Small but significant absolute ratio changes caused by the long odorant application**

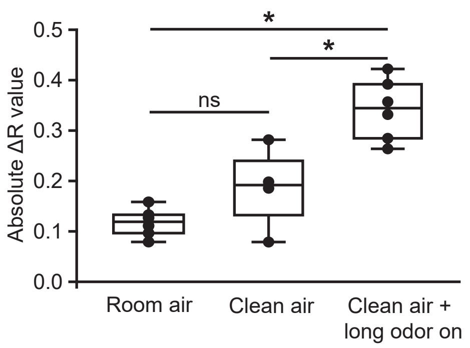

#### Fig. S6. Amyloid β deposition in the olfactory bulb of APPswe/PS1G384A mice

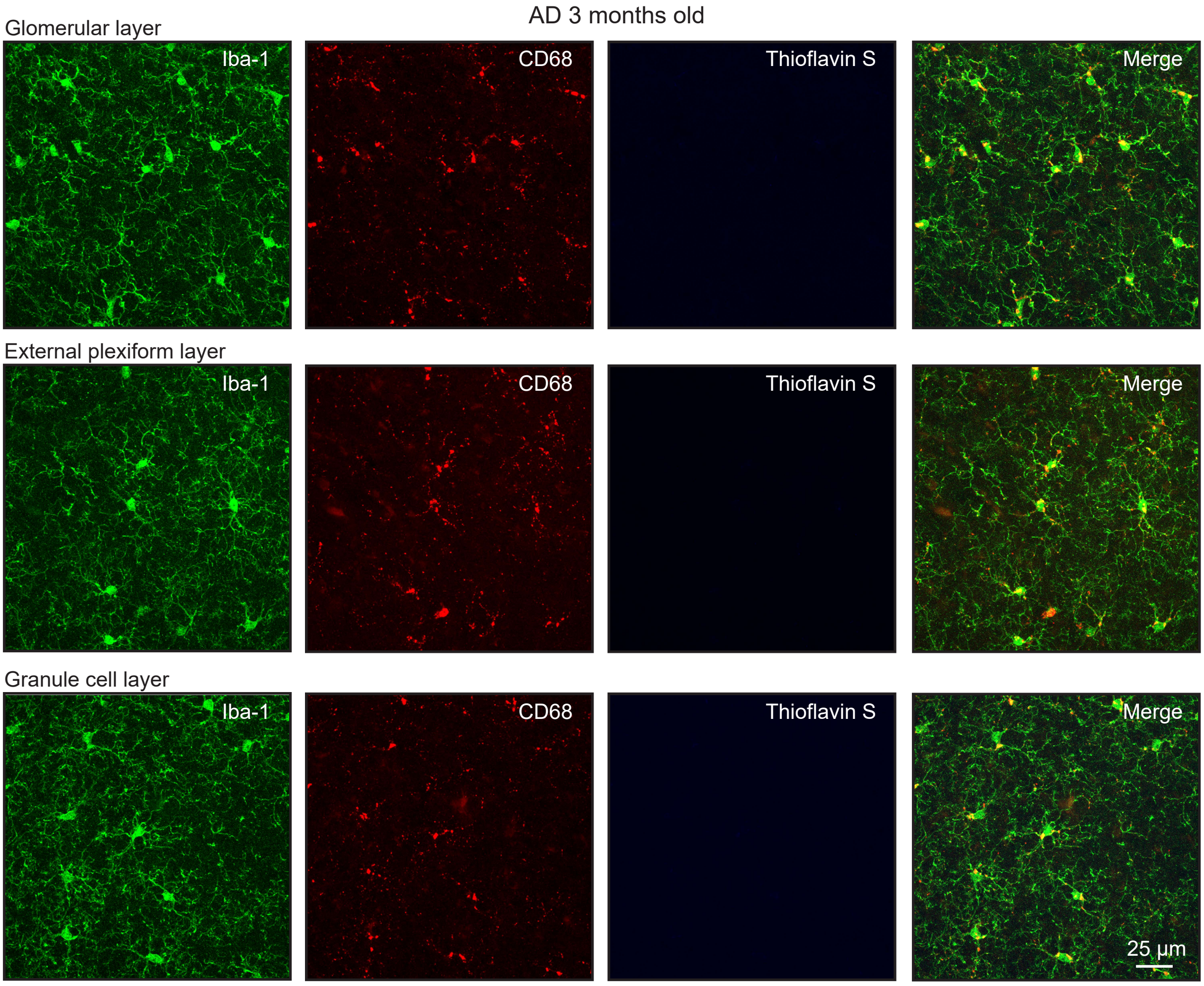


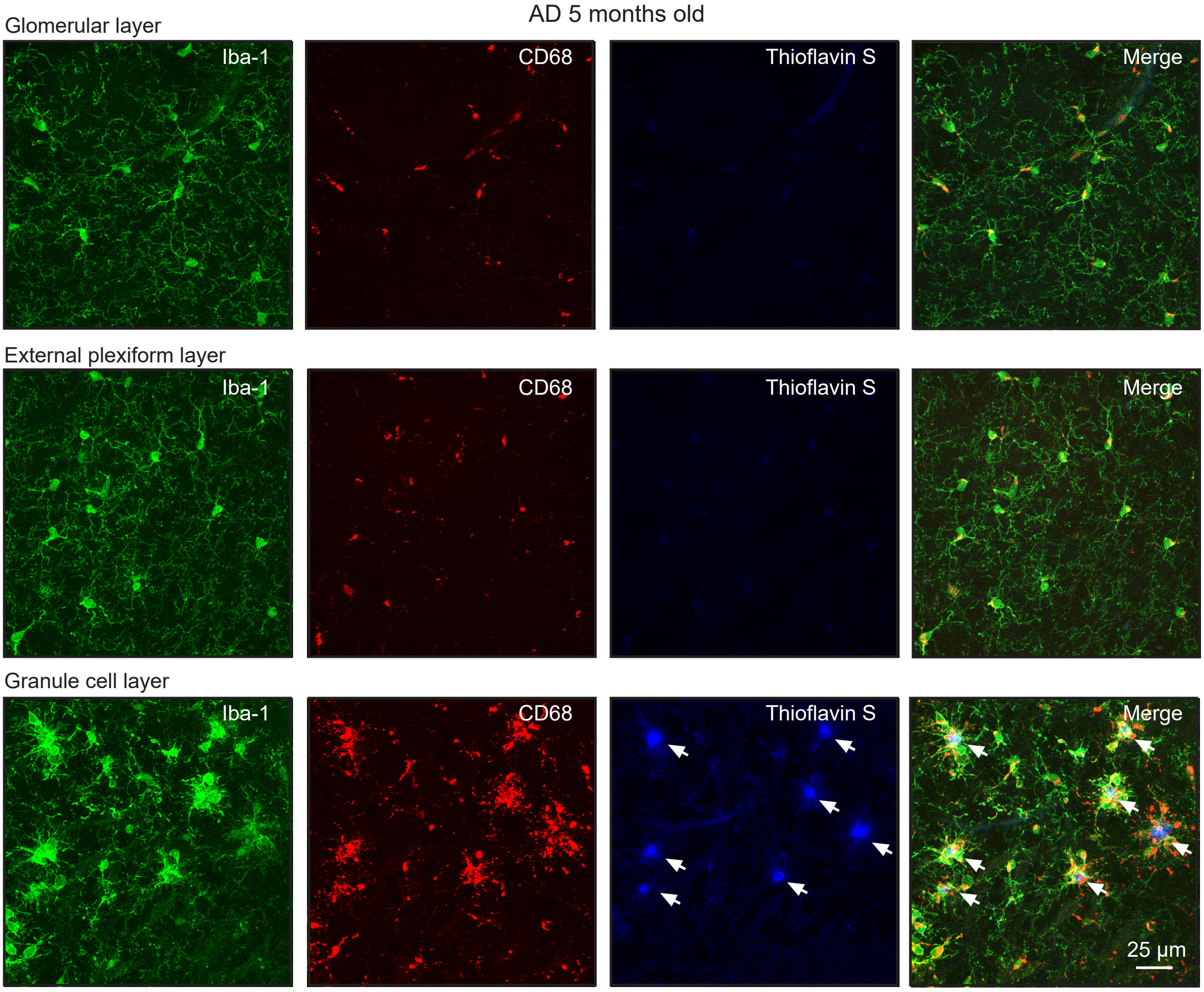

**Fig. S7. Little difference in the basal Twitch-2B ratios of mitral/tufted cells and their intercellular variance between WT and AD mice**

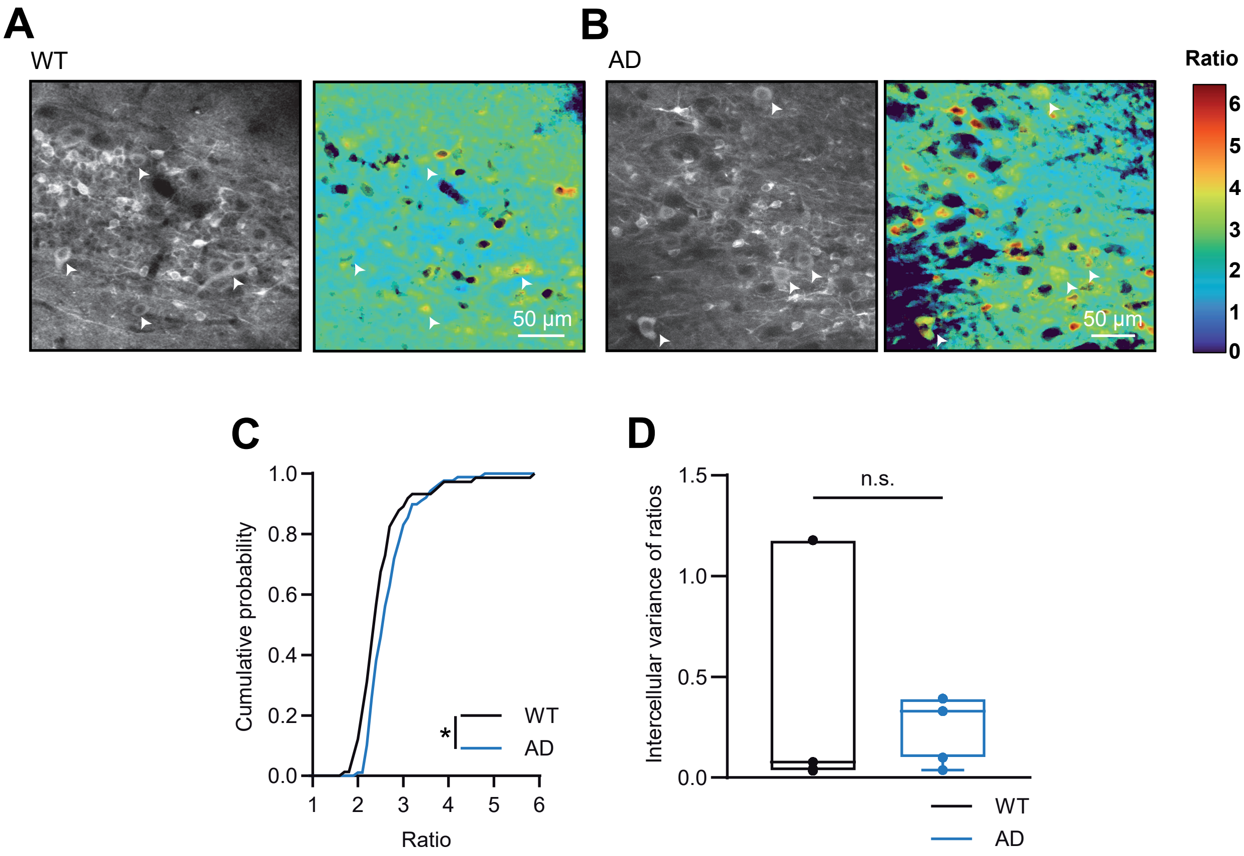

**Fig. S8. Presynaptic structures contribute to the glomerular Twitch-2B signal**

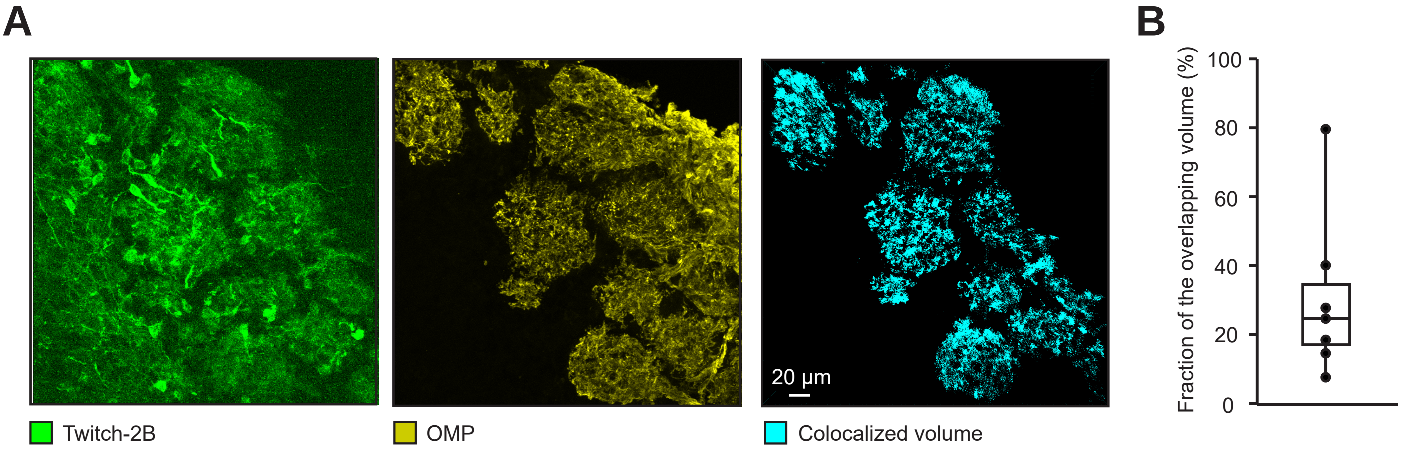
